# Genomics of Mesolithic Scandinavia reveal colonization routes and high-latitude adaptation

**DOI:** 10.1101/164400

**Authors:** Torsten Günther, Helena Malmström, Emma M. Svensson, Ayça Omrak, Federico Sánchez-Quinto, Gülşah M. Kılınç, Maja Krzewińska, Gunilla Eriksson, Magdalena Fraser, Hanna Edlund, Arielle R. Munters, Alexandra Coutinho, Luciana G. Simões, Mário Vicente, Anders Sjölander, Berit Jansen Sellevold, Roger Jørgensen, Peter Claes, Mark D. Shriver, Cristina Valdiosera, Mihai G. Netea, Jan Apel, Kerstin Lidén, Birgitte Skar, Jan Storå, Anders Götherström, Mattias Jakobsson

**Affiliations:** Department of Organismal Biology, Uppsala University, Norbyvägen 18C, 75236, Uppsala, Sweden; Department of Archaeology and Classical Studies, Stockholm University, Wallenberglaboratoriet, 10691 Stockholm, Sweden; Middle East Technical University, Department of Biological Sciences, 06800, Ankara, Turkey; Department of Archaeology and Ancient History, Uppsala University-Campus Gotland, 62167 Visby, Sweden; NIKU - Norwegian Institute for Cultural Heritage Research; Tromsø University Museum, UiT The Arctic University of Norway, P.O. Box 6050 Langnes, NO-9037 Tromsø, Norway; Department of Electrical Engineering, ESAT/PSI, KU Leuven, 3000, Leuven, Belgium; Department of Anthropology, Penn State University, 16801, University Park, USA; Department of Archaeology and History, La Trobe University, Melbourne, VIC 3086, Australia; Department of Internal Medicine and Radboud Center for Infectious Diseases, Radboud University Medical Center, Geert Grooteplein 8, 6525GA Nijmegen, The Netherlands; Department of Archaeology and Ancient History, Lund University, 22100, Lund, Sweden; Department of Archaeology and and Cultural History, NTNU University Museum, 7491 Trondheim, Norway; SciLife Lab

## Abstract

Scandinavia was one of the last geographic areas in Europe to become habitable for humans after the last glaciation. However, the origin(s) of the first colonizers and their migration routes remain unclear. We sequenced the genomes, up to 57x coverage, of seven hunter-gatherers excavated across Scandinavia and dated to 9,500-6,000 years before present. Surprisingly, among the Scandinavian Mesolithic individuals, the genetic data display an east-west genetic gradient that opposes the pattern seen in other parts of Mesolithic Europe. This result suggests that Scandinavia was initially colonized following two different routes: one from the south, the other from the northeast. The latter followed the ice-free Norwegian north Atlantic coast, along which novel and advanced pressure-blade stone-tool techniques may have spread. These two groups met and mixed in Scandinavia, creating a genetically diverse population, which shows patterns of genetic adaptation to high latitude environments. These adaptations include high frequencies of low pigmentation variants and a gene-region associated with physical performance, which shows strong continuity into modern-day northern Europeans.

## Introduction

As the ice-sheet retracted from northern Europe after the Last Glacial Maximum (LGM), around 23,000 years ago, new habitable areas emerged [1] allowing plants [2,3] and animals [4,5] to recolonize the Scandinavian peninsula (hereafter Scandinavia). There is consistent evidence of human presence in the archaeological record from c. 11,700 years before present (BP), both in southern and northern Scandinavia [6–9]. At this time, the ice-sheet was still dominating the interior of Scandinavia [9] (Fig. 1A, S1 Text), but recent climate modeling shows that the Arctic coast of (modern-day) northern Norway was ice-free [10]. Similarities in late-glacial lithic technology (direct blade percussion technique) of western Europe and the oldest counterparts of northernmost Scandinavia [11] (S1 Text) have been used to argue for a postglacial colonization of Scandinavia from southwestern Europe. However, studies of a new lithic technology, ‘pressure blade’ technique, which first occurred in the northern parts of Scandinavia, indicates contacts with groups in the east and possibly an eastern origin of the colonizers [7,12,13] (S1 Text). The first genetic studies of Mesolithic human remains from central and eastern Scandinavia (SHGs) revealed similarities to two different Mesolithic European populations, the ‘western hunter-gatherers’ (WHGs) from western, central and southern Europe and the ‘eastern hunter-gatherers’ (EHGs) from northeastern Europe [14–21]. Archaeology, climate modeling, and genetics, suggest several possibilities for the colonization of Scandinavia, including migrations from the south, southeast, northeast and combinations of these, however, the early post-glacial peopling of Scandinavia remains elusive [1,4,6–17,22,23]. In this study, we contrast genome sequence data and stable isotopes from Mesolithic human remains from western, northern, and eastern Scandinavia to infer the post-glacial colonization of Scandinavia – from where people came, what routes they followed, how they were related to other Mesolithic Europeans [15–19,24] – and to investigate human adaptation to high-latitude environments.

**Figure 1:**
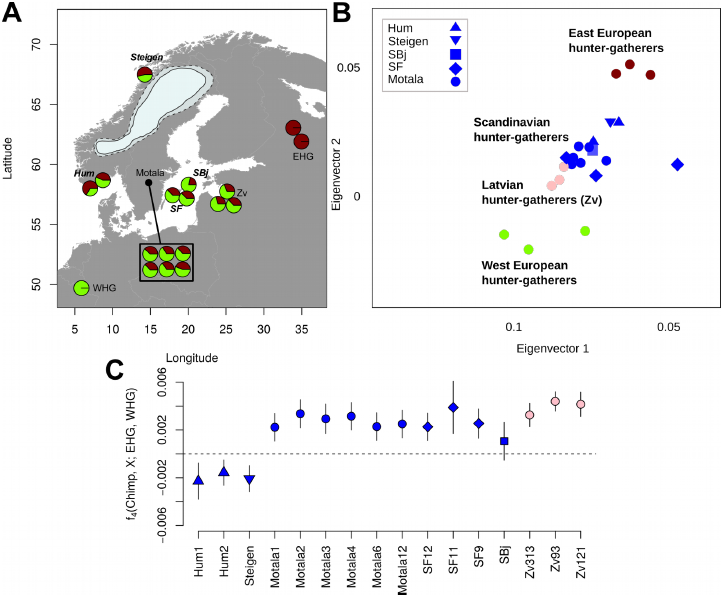
Mesolithic samples and their genetic affinities–. (A) Map of the Mesolithic European samples used in this study. The pie charts show the model-based [16,17] estimates of genetic ancestry for each SHG individual. The map also displays the ice sheet covering Scandinavia 10,000 BP (most credible (solid line) and maximum extend (dashed line) following [10]). Newly sequenced sites are shown in bold and italics, SF11 is excluded from this map due to its low coverage (0.1x). Additional European EHG and WHG individuals used in this study derive from sites outside this map. (B) Magnified section of genetic similarity among ancient and modern-day individuals using PCA featuring only the Mesolithic European samples (see S6 Text for the full plot). (C) Allele sharing between the SHGs, Latvian Mesolithic hunter-gatherers [33] and EHGs vs WHGs measured by f4(Chimpanzee, SHG; EHG, WHG) calculated for the captured SNPs for the EHGs [18]. Error bars show two block-jackknife standard errors.

## Results and Discussion

We sequenced the genomes of seven hunter-gatherers from Scandinavia (Table 1; S1 Text, S2 Text, S3 Text) ranging from 57.8× to 0.1× genome coverage, of which four individuals had a genome coverage above 1×. The remains were directly dated to between 9,500 BP and 6,000 BP, and were excavated in southwestern Norway (Hum1, Hum2), northern Norway (Steigen), and the Baltic islands of Stora Karlsö and Gotland (SF9, SF11, SF12 and SBj) and represent 18% (6 of 33) of all known human remains in Scandinavia older than 8,000 [25]. All samples displayed fragmentation and cytosine deamination at fragment termini characteristic for ancient DNA (S3 Text). Mitochondrial (mt) DNA-based contamination estimates were <6% for all individuals and autosomal contamination was <1% for all individuals except for SF11, which showed c. 10% contamination (Table 1, S4 Text). Four of the seven individuals were inferred to be males, three were females. All the western and northern Scandinavian individuals and one eastern Scandinavian carried U5a1 mitochondrial haplotypes while the remaining eastern Scandinavians carried U4a haplotypes (Table 1, S5 Text). These individuals represent the oldest U5a1 and U4 lineages detected so far. The Y chromosomal haplotype was determined for three of the four males, all carried I2 haplotypes, which were common in pre-Neolithic Europe (Table 1, S5 Text).

**Table 1:** Information on the seven Scandinavian hunter-gatherers investigated in this study, including calibrated date before present (cal BP) corrected for the marine reservoir effect, given as a range of two standard deviations, average genome coverage, average mitochondrial (mt) coverage, mt and Y chromosome haplogroups and contamination estimates based on the mt, the X-chromosome for males and the autosomes.

| Individual | Calibrated date (cal BP, 2 sigma) | Genome coverage | mt coverage | Sex | mt haplo-group | Y haplo-group | Contamination estimate |  |  |
| --- | --- | --- | --- | --- | --- | --- | --- | --- | --- |
|  |  |  |  |  |  |  | based on mt | based on X | based on autosomes |
| Hum1 | 9452-9275<sup>s</sup> | 0.71 | 597 | XX | U5a1 | - | 0.29% | - | 0.00% |
| Hum2 | 9452-9275<sup>s</sup> | 4.05 | 432 | XY | U5a1d | I2-L68 | 0.15% | 0.63% | 0.73% |
| Steigen | 5950-5764 | 1.24 | 277 | XY | U5a1d | I2a1b-M423 | 0.00% | 0.4% | 0.00% |
| SF9 | 9300-8988 | 1.15 | 93 | XX | U4a2 | - | 5.36% | - | 0.00% |
| SF11 | 9023-8760 | 0.10 | 45 | XY | U5a1 | * | 3.42% | * | 10.16% |
| SF12 | 9033-8757 | 57.79 | 9774 | XX | U4a1 | - | 0.34% | - | 0.932% |
| SBj | 8963-8579 | 0.43 | 102 | XY | U4a1 | I2-L68 | 3.72% | 1.4% | 0.06% |

<sup>s</sup> combined probability for the Hummervikholmen samples
\* not enough genome coverage

The high coverage and Uracil-DNA-glycosylase (UDG) treated genome (to reduce the effects of post-mortem DNA damage [26]) of SF12 allowed us to confidently discover new and hitherto unknown variants at sites with 55x or higher sequencing depth (S3 Text). Based on SF12’s high-coverage and high-quality genome, we estimate the number of single nucleotide polymorphisms (SNPs) hitherto unknown (that are not recorded in dbSNP (v142)) to be c. 10,600. This is almost twice the number of unique variants (c. 6,000) per Finnish individual (S3 Text) and close to the median per European individual in the 1000 Genomes Project [27] (c. 11,400, S3 Text). At least 17% of these SNPs that are not found in modern-day individuals, were in fact common among the Mesolithic Scandinavians (seen in the low coverage data conditional on the observation in SF12), suggesting that a substantial fraction of human variation has been lost in the past 9,000 years (S3 Text). In other words, the SHGs (as well as WHGs and EHGs) have no direct descendants, or a population that show direct continuity with the Mesolithic populations [14–17]. Thus, many genetic variants found in Mesolithic individuals have not been carried over to modern-day groups. Among the novel variants in SF12, four (all heterozygous) are predicted to affect the function of protein coding genes [28] (S3 Text). The ‘heat shock protein’ *HSPA2* in SF12 carries an unknown mutation that changes the amino acid histidine to tyrosine at a protein-protein interaction site, which likely disrupts the function of the protein (S3 Text). Defects in *HSPA2* are known to drastically reduce fertility in males [29]. Although SF12 herself would not be affected by this variant, her male offspring could carry the reduced fertility variant, and it will be interesting to see how common this variant was among Mesolithic groups as more genome sequence data become available. The high-quality diploid genotype calls further allowed us to genetically predict physical appearance, including pigmentation, and to use a model-based approach trained on modern-day faces and genotypes [30,31] to create a 3D model of SF12’s face (S9 Text). This represents a new way of reconstructing an ancient individual’s facial appearance from genetic information, which is especially informative in cases such as for SF12, where only post-cranial fragments were available, and future archaeogenetic studies will have the potential to many individuals appearance from past times.

### Demographic history of Mesolithic Scandinavians

In order to compare the genomic data of the seven SHGs to genetic information from other ancient individuals and modern-day groups, data was merged with six published Mesolithic individuals from Motala in central Scandinavia, 47 published Upper Paleolithic, Mesolithic and Early Neolithic individuals from other parts of Eurasia (S6 Text) [15–20,24,32–36], as well as with a world-wide set of 203 modern-day populations [16,27,37]. All 13 SHGs – regardless of geographic sampling location and age – display genetic affinities to both WHGs and EHGs (Fig. 1A, B, S6 Text). This is consistent with a scenario in which SHGs represent a mixed group tracing parts of their ancestry to both the WHGs and the EHGs [15–17,20,38].

To investigate the postglacial colonization of Scandinavia, we explored four hypothetical migration routes (primarily based on natural geography) linked to WHGs and EHGs, respectively (S11 Text); a) a migration of WHGs from the south, b) a migration of EHGs from the east across the Baltic Sea, c) a migration of EHGs from the east and along the north-Atlantic coast, d) a migration of EHGs from the east and south of the Baltic Sea, and combinations of these four migration routes. These scenarios allow us to formulate expected genetic affinities for northern, western, eastern, and central SHGs (S11 Text). The SHGs from northern and western Scandinavia show a distinct and significantly stronger affinity to the EHGs compared to the central and eastern SHGs (Fig. 1). Conversely, the SHGs from eastern and central Scandinavia were genetically more similar to WHGs compared to the northern and western SHGs (Fig. 1). Using [16,17], the EHG genetic component of northern and western SHGs was estimated to 55% on average (43-67%) and significantly different (Wilcoxon test, p=0.014) from the average 35% (22-44%) in eastern and south-central SHGs. This average is similar to eastern Baltic hunter-gatherers from Latvia [33] (average 33%, Fig. 1A, S6 Text). These patterns of genetic affinity within SHGs are in direct contrast to the expectation based on geographic proximity with EHGs and WHGs and do not correlate with age of the sample (S11 Text).

The archaeological record in Scandinavia shows early evidence of human presence in northern coastal Atlantic areas [13]. Stable isotope analysis of northern and western SHGs revealed an extreme marine diet, suggesting a maritime subsistence, in contrast to the more mixed terrestrial/aquatic diet of eastern and central SHGs (S1 Text). Combining these isotopic results with the patterns of genetic variation, we suggest an initial colonization from the south, likely by WHGs. A second migration of people who were related to the EHGs – that brought the new pressure blade technique to Scandinavia and that utilized the rich Atlantic coastal marine resources –entered from the northeast moving southwards along the ice-free Atlantic coast where they encountered WHG groups. The admixture between the two colonizing groups created the observed pattern of a substantial EHG component in the northern and the western SHGs, contrary to the higher levels of WHG genetic component in eastern and central SHGs (Fig. 1, S11 Text).

By sequencing complete ancient genomes, we can compute unbiased estimates of genetic diversity, which are informative of past population sizes and population history. Here, we restrict the analysis to WHGs and SHGs, since only SNP capture data is available for EHGs (S7 Text). In current-day Europe, there is greater genetic diversity in the south compared to the north. During the Mesolithic, by contrast, we find higher levels of genetic diversity (S7 Text) as well as lower levels of runs of homozygosity (Fig. 2A) and linkage disequilibrium (Fig. 2B) in SHGs compared to WHGs (represented by Loschbour and Bichon, [16,32]) and Caucasus hunter-gatherers (CHG, represented by Kotias and Satsurblia, [32]). Using a sequential-Markovian-coalescent approach [39] for the high-coverage, high quality genome of SF12, we find that right before the SF12 individual lived, the effective population size of SHGs was similar to that of WHGs (Fig. 2C). At the time of the LGM and back to c. 50,000 years ago, both the WHGs and SHGs go through a bottleneck, but the ancestors of SHGs retained a greater population size in contrast to the ancestors of WHGs who went through a more severe bottleneck (Fig. 2C). Around 50,000-70,000 years ago, the effective population sizes of the ancestors of SHGs, WHGs, Neolithic groups (represented by Stuttgart [16]) and Paleolithic Eurasians (represented by Ust-Ishim [36]) align, suggesting that these diverse groups all trace their ancestry back to a common ancestral group which likely represents the early migrants out-of-Africa, who likely share a common ancestry outside of Africa.

**Figure 2:**
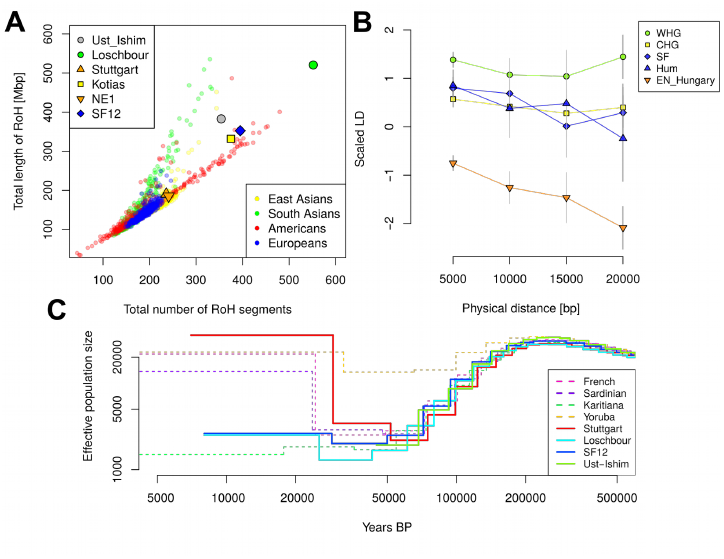
Genetic diversity in prehistoric Europe –. (A) Runs of Homozygosity (RoH) for the six prehistoric humans that have been sequenced to >20x genome coverage, (Kotias is a hunter-gatherer from the Caucasus region [32], NE1 is an early Neolithic individual from modern-day Hungary [24], the other individuals are described in the text), compared to all modern-day non-African individuals from the 1000 genomes project [27]. (B) Linkage disequilibrium (LD) decay for five prehistoric populations each represented by two individuals (eastern SHGs: SF (SF9 and SF12), western SHGs: Hum (Hum1 and Hum2), Caucasus hunter-gatherers [32]: CHG (Kotias and Satsurblia), WHGs [16,32] (Loschbour and Bichon), and early Neolithic Hungarians [24]: EN_Hungary (NE1 and NE6). LD was scaled in each distance bin by using the LD for two modern populations [27] as 1 (modern-day Tuscan, TSI) and as 0 (modern-day Peruvians, PEL). LD was calculated from the covariance of derived allele frequencies of two haploid individuals per population (S7 Text). Error bars show two standard errors estimate during 100 bootstraps across SNP pairs. (C) Effective population size over time as inferred by MSMC’ [39] for four prehistoric humans with high genome coverage. The dashed lines show the effective population sizes for modern-day populations. All curves for prehistoric individuals were shifted along the X axis according to their radiocarbon date.

### Adaptation to high-latitude environments

With the aim of detecting signs of adaptation to high-latitude environments and selection during and after the Mesolithic, we employed three different approaches that utilize the Mesolithic genomic data. In the first approach, we assumed that SHGs adapted to high-latitude environments of low temperatures and seasonally low levels of light, and searched for gene variants that carried over to modern-day people in northern Europe. As we have already noted, modern-day northern Europeans trace limited amount of genetic material back to the SHGs (due to the many additional migrations during later periods), and any genomic region that displays extraordinary genetic continuity would be a strong candidate for adaptation in people living in northern Europe across time. We designed a statistic, Dsel (S10 Text), that captures this specific signal and scanned the whole genome for gene-variants that show strong continuity (little differentiation) between SHGs and modern-day northern Europeans while exhibiting large differentiation to modern-day southern European populations [40] (Fig. 3A; S10 Text). Six of the top ten SNPs with greatest Dsel values were located in the *TMEM131* gene that has been found to be associated with physical performance [41], which could make it part of the physiological adaptation to cold [42]. This genomic region was more than 200kbp long and showed the strongest haplotypic differentiation between modern-day Tuscans and Finns (S10 Text). The particular haplotype was relatively common in SHGs, it is even more common among today’s Finnish population (S10 Text), and showed a strong signal of local adaptation (S10 Text). Other top hits included genes associated with a wide range of metabolic, cardiovascular, developmental and psychological traits (S10 Text) potentially linked to physiological [42].

**Figure 3:**
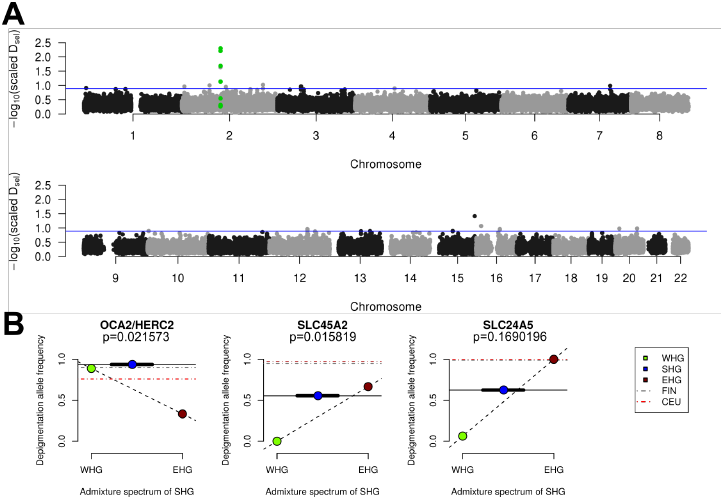
Adaptation to high-latitude climates –. (A) Manhattan plot of similarity between Mesolithic allele-frequency and modern-day Finnish (FIN) allele-frequency in contrast to difference to (TSI) allele-frequency using the statistic Dsel. The green-highlighted SNPs are all located in the *TMEM131* gene. The horizontal blue line depicts the top 0.01% Dsel SNPs across the genome. (B) Derived allele frequencies for three pigmentation associated SNPs (*SLC24A5, SLC45A2*, associated with skin pigmentation and *OCA2/HERC2* associated with eye pigmentation). The dashed line connecting EHG and WHG represents potential allele frequencies if SHG were a linear combination of admixture between EHG and WHG. The solid horizontal line represents the derived allele frequency in SHG. The blue symbols representing SHGs were set on the average genome-wide WHG/EHG mixture proportion (on x-axis) across all SHGs, the thick black line represents the minimum and maximum admixture proportions across all SHGs. Dashed horizontal lines represent modern European populations (CEU=Utah residents with Central European ancestry). The p-values were estimated from simulations of SHG allele frequencies based on their genome-wide ancestry proportions (S10 Text).

In addition to performing this genome-wide scan, we studied the allele frequencies in three pigmentation genes (*SLC24A5*, *SLC45A2*, having a strong effect on skin pigmentation, and *OCA2/HERC2*, having a strong effect on eye pigmentation) where the derived alleles are virtually fixed in northern Europeans today. The differences in allele frequencies of those three loci are among the highest between human populations, suggesting that selection was driving the differences in eye color, skin and hair pigmentation as part of the adaptation to different environments [43–46]. The SHGs show a combination of eye and skin pigmentation that was unique in Mesolithic Europe, with light skin pigmentation and varied blue to light-brown eye color. This is strikingly different from the WHGs – who have been found to have the specific combination of blue-eyes and dark-skin [16,18,19,21] (Fig. 3B) – and EHGs – who have been suggested to be brown eyed and light-skinned [17,18] (Fig. 3B). The unique configuration of the SHGs is not fully explained by the fact that SHGs are a mixture of EHGs and WHGs as the frequencies of the blue-eye and one light-skin variant are significantly higher in SHGs than expected from their genome-wide admixture proportions (Fig. 3B, S10 Text). This could be explained by a continued increase of the allele frequencies after the admixture event, likely caused by adaptation to high-latitude environments [43,45].

## Conclusion

By combining information from climate modeling, archaeology and Mesolithic human genomes, we were able to reveal the complexity of the early colonization process of Scandinavia and human adaptation to high-latitude environments. We disentangled migration routes and linked them to particular archaeological patterns, demonstrate greater genetic diversity in northern Europe compared to southern Europe – in contrast to modern-day patterns – and show that many genetic variants that were common in the Mesolithic have been lost today. These finds reiterate the importance of human migration for dispersal of novel technology in human prehistory [14–17,23,24,38,47–50] and the many partial population turnovers in our past.

## Materials and Methods

### Sample preparation

Genomic sequence data was generated from teeth and bone samples belonging to seven (eight, including SF13) Mesolithic Scandinavian hunter-gatherers (S1 Text). A detailed description on the archaeological background of the samples as well as post-LGM Scandinavia can be found in S1 Text. Additional libraries were sequenced for Ajvide58 and Ajvide70 [15] (S2 Text). All samples were prepared in dedicated ancient DNA (aDNA) facilities at the Evolutionary Biology Centre in Uppsala (SF9, SF11, SF12, SF13, SBj, Hum1, Hum2) and at the Archaeological research laboratory, Stockholm University (Steigen).

*DNA extraction and library building:* Bones and teeth were decontaminated prior to analysis by wiping them with a 1% Sodiumhypoclorite solution, and DNA free water. Further, all surfaces were UV irradiated (6 J/cm^2^ at 254 nm). After removing one millimeter of the surface, approximately 30-100 mg of bone was powderized and DNA was extracted following silica-based methods as in [51] with modifications as in [49,52] or as in [53] and eluted in 25-110 µl of EB buffer. Between one and 16 extractions were made from each sample and one extraction blank with water instead of bone powder was included per six to ten extracts. Blanks were carried along the whole process until qPCR and/or PCR and subsequent quantification.

DNA libraries were prepared using 20µl of extract, with blunt-end ligation coupled with P5 and P7 adapters and indexes as described in [49,54]. From each extract one to five double stranded libraries were built. Since aDNA is already fragmented the shearing step was omitted from the protocol. Library blank controls including water as well as extraction blanks were carried along during every step of library preparation. In order to determine the optimal number of PCR cycles for library amplification qPCR was performed. Each reaction was prepared in a total volume of 25 μl, containing 1 ul of DNA library, 1X MaximaSYBRGreen mastermix and 200 nM each of IS7 and IS8 [54] reactions were set up in duplicates. Each blunt-end library was amplified in four to 12 replicates with one negative PCR control per index-PCR. The amplification reactions had a total volume of 25 μl, with 3 ul DNA library, and the following in final concentrations; 1X AmpliTaq Gold Buffer, 2.5mM MgCl2, 250uM of each dNTP, 2.5U AmpliTaq Gold (Thermofisher), and 200nM each of the IS4 primer and index primer [54]. PCR was done with the following conditions; an activation step at 94°C for 10 min followed by 10-16 cycles of 94°C for 30s, 60°C for 30s and 72°C for 30s, and a final elongation step of 72°C for 10min. For each library four amplifications with the same indexing primer were pooled and purified with AMPure XP beads (Agencourt). The quality and quantity of libraries was checked using Tapestation or BioAnalyzer using the High Sensitivity Kit (Agilent Technologies). None of the blanks showed any presence of DNA comparable to that of a sample and were therefore not further analyzed. For initial screening 10-20 libraries were pooled at equimolar concentrations for sequencing on an Illumina HiSeq 2500 using v.4 chemistry and 125 bp paird-end reads or HiSeqX, 150bp paired-end length using v2.5 chemistry at the SNP & SEQ Technology Platform at Uppsala University. After evaluation of factors such as clonality, proportion of human DNA and genomic coverage samples were selected for re-sequencing aiming to yield as high coverage as possible for each library.

*Generation of a high coverage UDG treated genome:* Based on the results of the non-damage repair sequencing the SF12 individual was selected for large-scale sequencing in order to generate a high coverage genome of high quality where damages had been repaired using Uracil-DNA-glycosylase (UDG). In addition to the 15 extracts previously prepared and used for non-damage repair libraries, another 111 extracts were made based on a variety of silica based methods [24,49,51,52]. From these 126 extracts a total of 258 damage repaired double stranded libraries were built for Illumina sequencing platforms. Libraries were built as above, except a DNA repair step with (UDG and endonuclease VIII (endo VIII) or USER enzyme (NEB) treatment was included in order to remove deaminated cytosines [55]. Quantitative PCR (qPCR) was performed in order to quantify the number of molecules and the optimal number of PCR cycles prior to amplification for each DNA library. Furthermore, this step included extraction blanks, library blanks and amplification blanks to monitor potential contamination. All of these negative controls showed an optimal cycle of amplification significantly higher to those of our ancient DNA libraries (>10 cycles) and they were thus deemed as negative. Our experimental results show minimal levels of contamination, which is in concordance with mitochondrial DNA and X chromosome estimates of contamination (see S4 Text and Table 1). Each reaction was done in a total volume of 25 μl, containing 1 ul of DNA library, 1X MaximaSYBRGreen mastermix and 200nM each of IS7 and IS8 [54] reactions were set up in duplicate.The PCRs were set up using a similar system as for the non-damage repair samples (in quadruplicates that were pooled when the PCR products were cleaned), with the difference of using AccuPrime DNA polymerase instead of AmpliTaqGold (Thermofisher) and the following PCR conditions; an activation step at 95°C for 2 min followed by 10-16 cycles of 95°C for 15s, 60°C for 30s and 68°C for 1min, and a final elongation step of 68°C for 5min. Blank controls including water as well as extraction blanks were carried out during every step of library preparation. Amplified libraries were pooled, cleaned, quantified and sequenced in the same manner as non-damage repaired libraries. In order to sequence libraries to depletion, two to eight libraries were pooled together and sequenced until reaching a clonality of >50%, if the clonality was lower, the library was either classified as unproductive or when the sequencing goal (>55X coverage) was reached and further sequencing was deemed unnecessary. Sequencing was performed as above.

### Bioinformatic data processing and authentication

Paired-end reads were merged using MergeReadsFastQ_cc.py [56], if an overlap of at least 11 base pairs was found the base qualities were added together and any remaining adapters were trimmed. Merged reads were then mapped single ended with bwa aln 0.7.13 [57] to the human reference genome (build 36 and 37) using the following non-default parameters: seeds disabled -l 16500 -n 0.01 -o 2 [15,16]. To remove PCR duplicates, reads with identical start and end positions were collapsed using a modified version, to ensure random choice of bases, of FilterUniqSAMCons_cc.py [56]. Reads with less than 10 % mismatches to the human reference genome, reads longer than 35 base pairs and reads with mapping quality higher than 30 were used to estimate contamination.

The genetic data obtained from the two bone elements SF9 and SF13 showed extremely high similarities, which suggested that the two individuals were related. Using READ [58], a tool to estimate kin-relationship from ancient DNA, SF9 and SF13 were classified as either identical twins or the same individual. Therefore, we merged the genetic data for both individuals and refer to the merged individual as SF9 throughout the genetic analysis.

All data shows damage patterns indicative of authentic ancient DNA (S3 Text). Contamination was estimated using three different sources of data: (I) the mitochondrial genome [59], (II) the X chromosome if the individual was male [60,61] and (III) the autosomes[62]. Low contamination estimations over the three different approximations were interpreted as data mapping to the human genome being largely endogenous (S4 Text).

### Analysis of demographic history

Most population genomic analyses require a set of reference data for comparison. We compiled three different data sets from the literature and merged them with the data from ancient individuals (S6 Text). The three reference SNP panels were:
- The Human Origins genotype data set of 594,924 SNPs genotyped in 2,404 modern individuals from 203 populations [16,37].
- A panel of 1,055,209 autosomal SNPs which were captured in a set of ancient individuals by Mathieson et al [18].
- To reduce the potential effect of ascertainment bias on SNP array data and of cytosine deamination on transition SNPs, we also ascertained 1,797,398 transversion SNPs with a minor allele frequency of at least 10% (to avoid the effect of Eurasian admixture into Yorubans) in Yorubans of the 1000 genomes project [27]. Those SNPs were extracted using vcftools [63].

These data sets were merged with ancient individuals of less than 15x genome coverage using the following approach: for each SNP site, a random read covering that site with minimum mapping quality 30 was drawn (using samtools 0.1.19 mpileup [64]) and its allele was assumed to be homozygous in the ancient individual. Transition sites were coded as missing data for individuals that were not UDG treated and SNPs showing additional alleles or indels in the ancient individuals were excluded from the data.

Six high coverage ancient individuals (SF12, NE1 [24], Kotias [32], Loschbour [16], Stuttgart [16], Ust-Ishim [36]) used in this study were treated differently as we generated diploid genotype calls for them. First, the base qualities of all Ts in the first five base pairs of each read as well as all As in the last five base pairs were set to 2. We then used Picard [65] to add read groups to the files. Indel realignment was conducted with GATK 3.5.0 [66] using indels identified in phase 1 of the 1000 genomes project as reference [27]. Finally, GATK’s UnifiedGenotyper was used to call diploid genotypes with the parameters -stand_call_conf 50.0, -stand_emit_conf 50.0, -mbq 30, -contamination 0.02 and –output_mode EMIT_ALL_SITES using dbSNP version 142 as known SNPs. SNP sites from the reference data sets were extracted from the VCF files using vcftools [63] if they were not marked as low quality calls. Plink 1.9 [67,68] was used to merge the different data sets.

We performed principal component analysis (PCA) to characterize the genetic affinities of the ancient Scandinavian genomes to previously published ancient and modern genetic variation. PCA was conducted on 42 present-day west Eurasian populations from the Human Origins dataset [16,37], using *smartpca [69]* with numoutlieriter: 0 and lsqproject: YES options. A total of 59 ancient genomes (52 previously published and 7 reported here) (Table S6.1) were projected into the reference PCA space, computed from the genotype of modern individuals. For all individuals, a single allele was selected randomly making the data set fully homozygous. The result was plotted using the *ploteig* program of the EIGENSOFT [69] using with the –x and –k options.

*D and f statistics:* The qpDstat program of ADMIXTOOLS was used to calculate *D*-statistics to test deviations from a tree-like population topology of the shape ((*A*,*B*);(*X*,*Y*)) [37]. Standard errors were calculated using a block jackknife of 0.5 Mbp. The tree topologies are balanced at zero, indicating no recent interactions between the test populations. Significant deviations from zero indicate a deviation from the proposed tree topology depending on the value. Positive values indicate an excess of shared alleles between A and X or B and Y while negative values indicate more shared alleles between B and X or A and Y. Using an outgroup as population A limits the test results to depend on the recent relationships between B and Y (if positive) or B and X (if negative). Here we used high-coverage Mota [35], Yoruba [27] and Chimp genome as (A) outgroups. The software popstats [70] was used to calculate f4 statistics, in order to estimate shared drift between groups. Standard errors and Z scores for f4 statistics were estimated using a weighted block jackknife (Fig. 1C).

*Model-based clustering:* A model-based clustering algorithm, implemented in the ADMIXTURE software [71], was used to estimate ancestry components and to cluster individuals.

ADMIXTURE was conducted on the Human Origins data set [16,37], which was merged with the ancient individuals as described above. Data was pseudo-haploidized by randomly selecting one allele at each heterozygous site of present-day individuals. Finally, the dataset was filtered for linkage disequilibrium using PLINK [67,68] with parameters (–indep-pairwise 200 25 0.4), this retained 289,504 SNPs. ADMIXTURE was run in 50 replicates with different random seeds for ancestral clusters from K=2 to K=20. Common signals between independent runs for each K were identified using the LargeKGreedy algorithm of CLUMPP [72]. Clustering was visualized using rworldmap, ggplot2, SDMTools and RColorBrewer packages of GNU R version 3.3.0.

Starting from K=3, when the modern samples split up into an African and Eastern and Western Eurasian clusters, the Mesolithic Scandinavians from Norway show slightly higher proportions of the Eastern cluster than Swedish Mesolithic individuals. This pattern continues to develop across higher values of K and it is consistent with the higher Eastern affinities of the Norwegian samples seen in the PCA and D/f4 statistics. The results for all Ks are shown in S1 Fig.

In addition to ADMIXTURE, we assessed the admixture patterns in Mesolithic Scandinavians using a set of methods implemented in ADMIXTOOLS [37], qpWave [73] and qpAdm [16,17]. Both methods are based on f4 statistics, which relate a set of test populations to a set of outgroups in different distances from the potential source populations. We used the following set of outgroup populations from the Human Origins data set: Ami_Coriell, Biaka, Bougainville, Chukchi, Eskimo_Naukan, Han, Karitiana, Kharia, Onge. We first used qpWave to test the number of source populations for Mesolithic West Eurasians (WHG). qpWave calculates a set of statistics X(u,v) = f4(u0, u; v0, v) where u0 and v0 are populations from the sets of test populations L and outgroups R, respectively. To avoid having more test populations than outgroups, we built four groups consisting of (a) genetically western and central hunter-gatherers (Bichon, Loschbour, KO1, LaBrana), (b) Eastern hunter-gatherers (UzOO74/I0061, SVP44/I0124, UzOO40/I0211) (c) Norwegian hunter-gathers (Hum1, Hum2, Steigen) and (d) Swedish hunter-gatherers (individuals from Motala and Mesolithic Gotland). qpWave tests the rank of the matrix of all X(u,v) statistics. If the matrix has rank m, the test populations can be assumed to be related to at least m+1 “waves” of ancestry, which are differently related to the outgroups. A rank of 0 is rejected in our case (p=3.13e-81) while a rank of 1 is consistent with the data (p=0.699). Haak et al 2015 already showed, using the same approach, that WHG and EHG descend from at least two sources (confirmed with our data as rank 0 is rejected with p=1.66e-86, while rank 1 is consistent with the data) and adding individuals from Motala does not change these observations. Therefore, we conclude that European Mesolithic populations, including Swedish and Norwegian Mesolithic individuals, have at least two source populations.

We then used qpAdm to model Mesolithic Scandinavian individuals as a 2-way admixture of WHG and EHG. qpAdm was run separately for each Scandinavian individual x, setting T=x as target and S={EHG, WHG} as sources. The general approach of qpAdm is related to qpWave: target and source are used as L (with T being the base population) and f4 statistics with outgroups from R (same as above) are calculated. The rank of the resulting matrix is then set to the number of sources minus one, which allows to estimate the admixture contributions from each populations in S to T. The results are shown in Fig. 1.

*Runs of homozygosity:* Heterozygosity is a measurement for general population diversity and its effective population size. Analyzing the extent of homozygous segments across the genome can also give us a temporal perspective on the effective population sizes. Many short segments of homozygous SNPs can be connected to historically small population sizes while an excess of long runs of homozygosity suggests recent inbreeding. We restricted this analysis to the six high coverage individuals (SF12, NE1, Kotias, Loschbour, Stuttgart, Ust-Ishim) for which we obtained diploid genotype calls and we compared them to modern individuals from the 1000 genomes project. The length and number of runs of homozygosity were estimated using Plink 1.9 [67,68] and the parameters –homozyg-density 50, –homozyg-gap 100, –homozyg-kb 500, –homozyg-snp 100, –homozyg-window-het 1, –homozyg-window-snp 100, –homozyg-window-threshold 0.05 and –homozyg-window-missing 20. The results are shown in Fig. 2A.

*Linkage disequilibrium:* Similar to runs of homozygosity, the decay of linkage disequilibrium (LD) harbors information on the demographic history of a population. Long distance LD can be caused by a low effective population size and past bottlenecks. Calculating LD for ancient DNA data is challenging as the low amounts of authentic DNA usually just yields haploid allele calls with unknown phase. In order to estimate LD decay for ancient populations we first combine two haploid ancient individuals to a pseudo-diploid individual (similar to the approach chosen for conditional nucleotide diversity, S7 Text). Next, we bin SNP pairs by distance (bin size 5kb) and then calculate the covariance of derived allele frequencies (0, 0.5 or 1.0) for each bin. This way, we do not need phase information to calculate LD decay as we do not consider multilocus haplotypes, which is similar to the approach taken by ROLLOFF[37,74] and ALDER [75] to date admixture events based on admixture LD decay. For Fig. 2B, we used two modern 1000 genomes populations to scale the LD per bin. The LD between two randomly chosen PEL (Peruvian) individuals was set to 1 and the LD between two randomly chosen TSI (Tuscan) individuals was set to 0. This approach is used to obtain a relative scale for the ancient populations and we caution against a direct interpretation of the differences to modern populations as technical differences in the modern data (e.g. SNP calling or imputation) may have substantial effects.

*Effective population size:* We are using MSMC’s implementation of PSMC’ [39] to infer effective population sizes over time from single high coverage genomes. We restrict this analysis to UDG-treated individuals (SF12, Loschbour, Stuttgart, Ust-Ishim) as post-mortem damage would cause an excess of false heterozygous transition sites. Input files were prepared using scripts provided with the release of MSMC (https://github.com/stschiff/msmc-tools) and MSMC was run with the non-default parameters –fixedRecombination and -r 0.88 in order to set the ratio of recombination to mutation rate to a realistic level for humans. We also estimate effective population size for six high-coverage modern genomes [76] (Fig. 2C). We plot the effective population size assuming a mutation rate of 1.25x10e-8 and a generation time of 30 years. The curves for ancient individuals were shifted based on their average C14 date.

### Detecting adaptation to high-latitude environments

We scanned the genomes for SNPs with similar allele frequencies in Mesolithic and modern-day northern Europeans, and contrast it to a modern-day population from southern latitudes. Pooling all Mesolithic Scandinavians together, we obtain an allele frequency estimate for Scandinavian hunter-gatherers (SHG) which is compared to modern-day Finnish individuals (FIN) and Tuscan individuals (TSI) from the 1000 genomes project [27]. We use the Finnish population as representatives of modern-day northern Europeans (this sample contains the largest number of sequenced genomes from a northern European population). Tuscans are used as an alternative population, who also traces some ancestry to Mesolithic populations, but who do not trace their ancestry to groups that lived at northern latitudes in the last 7-9,000 years. Our approach is similar to PBS [77] and inspired by DAnc [40], for each SNP, we calculate the statistic Dsel comparing the allele frequencies between an ancestral and two modern populations:

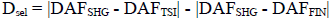

This scan was performed on all transversion SNPs extracted from the 1000 genomes data. Only sites with a high confidence ancestral allele in the human ancestor (as used by the 1000 genomes project [27]) and with coverage for at least six ancient Scandinavians were included in the com-putation. More information can be found in S10 Text.

## Acknowledgements

We thank Rachel Howcroft, and Nicci Arosén for help with isotope laboratory work and Heike Siegmund at the Stable Isotope Laboratory, Stockholm University, Lena Ideström at Gotlands museum and Sabine Sten at Uppsala University Campus Gotland in assisting in sampling the Stora Bjers material, Leena Drenzel at The Swedish History Museum for assistance with the Stora Förvar material, and the Norwegian Maritime Museum for excavating Hummervikholmen. MS and PC thank the participants who volunteered for face reconstruction methods development. This project was supported by grants from Riksbankens Jubileumsfond (to AG, MJ, and JS), Knut and Alice Wallenberg foundation (to MJ, JS, AG), the Swedish Research council, no. 2013-1905 (to AG, MJ, and JS) and no. 421-2013-730 (to JA and JS), an European Research Council Start-ing Grant (to MJ), a Wenner-Gren foundations postdoctoral fellowship (to TG), and Berit Wallen-berg foundation (to MF). Sequencing was performed at the National Genomics Infrastructure (NGI) Uppsala and computations were performed at Uppsala Multidisciplinary Center for Ad-vanced Computational Science (UPPMAX).

## References

Andersen B , Borns HW. The ice age world: an introduction to quaternary history and research with emphasis on North America and Northern Europe during the last 2.5 million years. Oslo [u.a.: Scandinavian Univ. Press; 1994.

Parducci L , Jorgensen T , Tollefsrud MM , Elverland E , Alm T , Fontana SL , et al. Glacial Survival of Boreal Trees in Northern Scandinavia. Science. 2012;335: 1083–1086. doi:10.1126/science.1216043

François O , Blum MGB , Jakobsson M , Rosenberg NA. Demographic History of European Populations of Arabidopsis thaliana. PLOS Genetics. 2008;4: e1000075. doi:10.1371/journal.pgen.1000075

Hewitt G. The genetic legacy of the Quaternary ice ages. Nature. 2000;405: 907–913. doi:10.1038/35016000

Xenikoudakis G , Ersmark E , Tison J-L , Waits L , Kindberg J , Swenson JE , et al. Consequences of a demographic bottleneck on genetic structure and variation in the Scandinavian brown bear. Mol Ecol. 2015;24: 3441–3454. doi:10.1111/mec.13239

Riede F. The resettlement of Northern Europe. In: Cummings V , Jordan P , Zvelebil M , editors. The Oxford Handbook of the Archaeology and Anthropology of Hunter-Gatherers. Oxford University Press; 2014.

Kankaanpää J , Rankama T. Fast or slow pioneers? A view from northern Lapland. In: Riede F , Tallavaara M , editors. Lateglacial and Postglacial Pioneers in Northern Europe, BAR International Series. Oxford: Oxford Archaeopress; 2014. pp. 147–154.

Breivik HM. Palaeo-oceanographic development and human adaptive strategies in the Pleistocene– Holocene transition: A study from the Norwegian coast. The Holocene. 2014;24: 1478–1490.

Stroeven AP , Hättestrand C , Kleman J , Heyman J , Fabel D , Fredin O , et al. Deglaciation of Fennoscandia. Quaternary Science Reviews. 2016;147: 91–121.

Hughes AL , Gyllencreutz R , Lohne ØS , Mangerud J , Svendsen JI. The last Eurasian ice sheets–a chronological database and time-slice reconstruction, DATED-1. Boreas. 2016;45: 1–45.

Bjerck HB , Breivik HM , Piana EL , Zangrando AF. Exploring the role of pinnipeds in the human colonization of the seascapes of Patagonia and Scandinavia. In: Bjerck HB , Breivik HM , Fretheim SE , Piana EL , Skar B , Tivoli AM , et al., editors. Equinox Sheffield; 2016. pp. 53–74.

Sørensen M , Rankama T , Kankaanpää J , Knutsson K , Knutsson H , Melvold S , et al. The First Eastern Migrations of People and Knowledge into Scandinavia: Evidence from Studies of Mesolithic Technology, 9th-8th Millennium BC. Norwegian Archaeological Review. 2013;46: 19–56. doi:10.1080/00293652.2013.770416

Knutsson H , Knutsson K , Molin F , Zetterlund P. From flint to quartz: Organization of lithic technology in relation to raw material availability during the pioneer process of Scandinavia. Quaternary International. 2016;424: 32–57. doi:10.1016/j.quaint.2015.10.062

Skoglund P , Malmstrom H , Raghavan M , Stora J , Hall P , Willerslev E , et al. Origins and Genetic Legacy of Neolithic Farmers and Hunter-Gatherers in Europe. Science. 2012;336: 466–469. doi:10.1126/science.1216304

Skoglund P , Malmstrom H , Omrak A , Raghavan M , Valdiosera C , Gunther T , et al. Genomic Diversity and Admixture Differs for Stone-Age Scandinavian Foragers and Farmers. Science. 2014;344: 747–750. doi:10.1126/science.1253448

Lazaridis I , Patterson N , Mittnik A , Renaud G , Mallick S , Kirsanow K , et al. Ancient human genomes suggest three ancestral populations for present-day Europeans. Nature. 2014;513: 409–413. doi:10.1038/nature13673

Haak W , Lazaridis I , Patterson N , Rohland N , Mallick S , Llamas B , et al. Massive migration from the steppe was a source for Indo-European languages in Europe. Nature. 2015;522: 207–211. doi:10.1038/nature14317

Mathieson I , Lazaridis I , Rohland N , Mallick S , Patterson N , Roodenberg SA , et al. Genome-wide patterns of selection in 230 ancient Eurasians. Nature. 2015;528: 499–503. doi:10.1038/nature16152

Olalde I , Allentoft ME , Sánchez-Quinto F , Santpere G , Chiang CWK , DeGiorgio M , et al. Derived immune and ancestral pigmentation alleles in a 7,000-year-old Mesolithic European. Nature. 2014;507: 225–228. doi:10.1038/nature12960

Raghavan M , Skoglund P , Graf KE , Metspalu M , Albrechtsen A , Moltke I , et al. Upper Palaeolithic Siberian genome reveals dual ancestry of Native Americans. Nature. 2013;505: 87–91. doi:10.1038/nature12736

Fu Q , Posth C , Hajdinjak M , Petr M , Mallick S , Fernandes D , et al. The genetic history of Ice Age Europe. Nature. 2016;534: 200–5. doi:10.1038/nature17993

Cummings V , Jordan P , Zvelebil M , editors. The Oxford Handbook of the Archaeology and Anthropology of Hunter-Gatherers. Oxford University Press; 2014. doi:10.1093/oxfordhb/9780199551224.001.0001

Allentoft ME , Sikora M , Sjögren K-G , Rasmussen S , Rasmussen M , Stenderup J , et al. Population genomics of Bronze Age Eurasia. Nature. 2015;522: 167–172. doi:10.1038/nature14507

Gamba C , Jones ER , Teasdale MD , McLaughlin RL , Gonzalez-Fortes G , Mattiangeli V , et al. Genome flux and stasis in a five millennium transect of European prehistory. Nature Communications. 2014;5: 5257. doi:10.1038/ncomms6257

Sjögren K-G , Ahlström T. Early Mesolithic burials from Bohuslän, western Sweden. Mesolithic burials – Rites, symbols and social organisation of early postglacial communities. Beier & Beran; 2017.

Briggs AW , Stenzel U , Meyer M , Krause J , Kircher M , Pääbo S. Removal of deaminated cytosines and detection of in vivo methylation in ancient DNA. Nucleic Acids Res. 2010;38: e87–e87. doi:10.1093/nar/gkp1163

Auton A , Abecasis GR , Altshuler DM , Durbin RM , Abecasis GR , Bentley DR , et al. A global reference for human genetic variation. Nature. 2015;526: 68–74. doi:10.1038/nature15393

Cingolani P , Platts A , Wang LL , Coon M , Nguyen T , Wang L , et al. A program for annotating and predicting the effects of single nucleotide polymorphisms, SnpEff: SNPs in the genome of Drosophila melanogaster strain w ^1118^; iso-2; iso-3. Fly. 2012;6: 80–92. doi:10.4161/fly.19695

Nixon B , Bromfield E , Dun M , Redgrove K , McLaughlin E , Aitken Rj. The role of the molecular chaperone heat shock protein A2 (HSPA2) in regulating human sperm-egg recognition. Asian Journal of Andrology. 2015;17: 568. doi:10.4103/1008-682X.151395

Claes P , Liberton DK , Daniels K , Rosana KM , Quillen EE , Pearson LN , et al. Modeling 3D Facial Shape from DNA. Luquetti D , editor. PLoS Genetics. 2014;10: e1004224. doi:10.1371/journal.pgen.1004224

Claes P , Hill H , Shriver MD. Toward DNA-based facial composites: preliminary results and validation. Forensic Science International: Genetics. 2014;13: 208–216.

Jones ER , Gonzalez-Fortes G , Connell S , Siska V , Eriksson A , Martiniano R , et al. Upper Palaeolithic genomes reveal deep roots of modern Eurasians. Nature communications. 2015;6: 8912. doi:10.1038/ncomms9912

Jones ER , Zarina G , Moiseyev V , Lightfoot E , Nigst PR , Manica A , et al. The Neolithic Transition in the Baltic Was Not Driven by Admixture with Early European Farmers. Curr Biol. 2017;27: 576–582. doi:10.1016/j.cub.2016.12.060

Seguin-Orlando A , Korneliussen TS , Sikora M , Malaspinas A-S , Manica A , Moltke I , et al. Genomic structure in Europeans dating back at least 36,200 years. Science. 2014;346: 1113–1118. doi:10.1126/science.aaa0114

Llorente MG , Jones ER , Eriksson A , Siska V , Arthur KW , Arthur JW , et al. Ancient Ethiopian genome reveals extensive Eurasian admixture throughout the African continent. Science. 2015;350: 820–2. doi:10.1126/science.aad2879

Fu Q , Li H , Moorjani P , Jay F , Slepchenko SM , Bondarev AA , et al. Genome sequence of a 45,000-year-old modern human from western Siberia. Nature. 2014;514: 445–449. doi:10.1038/nature13810

Patterson N , Moorjani P , Luo Y , Mallick S , Rohland N , Zhan Y , et al. Ancient Admixture in Human History. Genetics. 2012;192: 1065–1093. doi:10.1534/genetics.112.145037

Günther T , Jakobsson M. Genes mirror migrations and cultures in prehistoric Europe-a population genomic perspective. Curr Opin Genet Dev. 2016;41: 115–123. doi:10.1016/j.gde.2016.09.004

Schiffels S , Durbin R. Inferring human population size and separation history from multiple genome sequences. Nature genetics. 2014;46: 919–25. doi:10.1038/ng.3015

Key FM , Fu Q , Romagné F , Lachmann M , Andrés AM. Human adaptation and population differentiation in the light of ancient genomes. Nature Communications. 2016;7: 10775. doi:10.1038/ncomms10775

Vasan RS , Larson MG , Aragam J , Wang TJ , Mitchell GF , Kathiresan S , et al. Genome-wide association of echocardiographic dimensions, brachial artery endothelial function and treadmill exercise responses in the Framingham Heart Study. BMC Medical Genetics. 2007;8: S2. doi:10.1186/1471-2350-8-S1-S2

Makinen TM. Different types of cold adaptation in humans. Frontiers in bioscience. 2010;2: 1047–1067. doi:10.2741/S117

Fan S , Hansen MEB , Lo Y , Tishkoff SA. Going global by adapting local: A review of recent human adaptation. Science. 2016;354: 54–59. doi:10.1126/science.aaf5098

Donnelly MP , Paschou P , Grigorenko E , Gurwitz D , Barta C , Lu RB , et al. A global view of the OCA2-HERC2 region and pigmentation. Human Genetics. 2012;131: 683–696. doi:10.1007/s00439-011-1110-x

Jablonski NG , Chaplin G. The colours of humanity: the evolution of pigmentation in the human lineage. Philosophical Transactions of the Royal Society B: Biological Sciences. 2017;372: 20160349. doi:10.1098/rstb.2016.0349

Nielsen R , Akey JM , Jakobsson M , Pritchard JK , Tishkoff S , Willerslev E. Tracing the peopling of the world through genomics. Nature. 2017;541: 302–310. doi:10.1038/nature21347

Hofmanová Z , Kreutzer S , Hellenthal G , Sell C , Diekmann Y , Díez-del-Molino D , et al. Early farmers from across Europe directly descended from Neolithic Aegeans. PNAS. 2016; 201523951. doi:10.1073/pnas.1523951113

Cassidy LM , Martiniano R , Murphy EM , Teasdale MD , Mallory J , Hartwell B , et al. Neolithic and Bronze Age migration to Ireland and establishment of the insular Atlantic genome. Proceedings of the National Academy of Sciences. 2015; 1–6. doi:10.1073/pnas.1518445113

Günther T , Valdiosera C , Malmström H , Ureña I , Rodriguez-Varela R , Sverrisdóttir ÓO , et al. Ancient genomes link early farmers from Atapuerca in Spain to modern-day Basques. Proc Natl Acad Sci USA. 2015;112: 11917–11922. doi:10.1073/pnas.1509851112

Olalde I , Schroeder H , Sandoval-Velasco M , Vinner L , Lobón I , Ramirez O , et al. A common genetic origin for early farmers from mediterranean cardial and central european LBK cultures. Molecular Biology and Evolution. 2015;32: 3132–3142. doi:10.1093/molbev/msv181

Yang DY , Eng B , Waye JS , Dudar JC , Saunders SR. Technical note: improved DNA extraction from ancient bones using silica-based spin columns. Am J Phys Anthropol. 1998;105: 539–543. doi:10.1002/(SICI)1096-8644(199804)105:4<539::AID-AJPA10>3.0.CO;2-1

Malmström H , Svensson EM , Gilbert MTP , Willerslev E , Götherström A , Holmlund G. More on contamination: The use of asymmetric molecular behavior to identify authentic ancient human DNA. Molecular Biology and Evolution. 2007;24: 998–1004. doi:10.1093/molbev/msm015

Dabney J , Knapp M , Glocke I , Gansauge M-T , Weihmann A , Nickel B , et al. Complete mitochondrial genome sequence of a Middle Pleistocene cave bear reconstructed from ultrashort DNA fragments. Proc Natl Acad Sci USA. 2013;110: 15758–15763. doi:10.1073/pnas.1314445110

Meyer M , Kircher M. Illumina Sequencing Library Preparation for Highly Multiplexed Target Capture and Sequencing. Cold Spring Harbor Protocols. 2010;2010: pdb.prot5448–pdb.prot5448. doi:10.1101/pdb.prot5448

Briggs AW , Heyn P. Preparation of next-generation sequencing libraries from damaged DNA. Methods in Molecular Biology. 2012;840: 143–154. doi:10.1007/978-1-61779-516-9_18

Kircher M. Analysis of high-throughput ancient DNA sequencing data. Methods Mol Biol. 2012;840: 197–228. doi:10.1007/978-1-61779-516-9_23

Li H , Durbin R. Fast and accurate short read alignment with Burrows-Wheeler transform. Bioinformatics. 2009;25: 1754–1760. doi:10.1093/bioinformatics/btp324

Kuhn JMM , Jakobsson M , Günther T. Estimating genetic kin relationships in prehistoric populations. bioRxiv. 2017; 100297.

Green RE , Malaspinas A-S , Krause J , Briggs AW , Johnson PLF , Uhler C , et al. A Complete Neandertal Mitochondrial Genome Sequence Determined by High-Throughput Sequencing. Cell. 2008;134: 416–426. doi:10.1016/j.cell.2008.06.021

Rasmussen M , Guo X , Wang Y , Lohmueller KE , Rasmussen S , Albrechtsen A , et al. An Aboriginal Australian genome reveals separate human dispersals into Asia. Science. 2011;334: 94–98. doi:10.1126/science.1211177

Korneliussen TS , Albrechtsen A , Nielsen R. ANGSD: Analysis of Next Generation Sequencing Data. BMC Bioinformatics. 2014;15: 356. doi:10.1186/s12859-014-0356-4

Jun G , Flickinger M , Hetrick KN , Romm JM , Doheny KF , Abecasis GR , et al. Detecting and estimating contamination of human DNA samples in sequencing and array-based genotype data. American Journal of Human Genetics. 2012;91: 839–848. doi:10.1016/j.ajhg.2012.09.004

Danecek P , Auton A , Abecasis G , Albers CA , Banks E , DePristo MA , et al. The variant call format and VCFtools. Bioinformatics. 2011;27: 2156–2158. doi:10.1093/bioinformatics/btr330

Li H , Handsaker B , Wysoker A , Fennell T , Ruan J , Homer N , et al. The Sequence Alignment/Map format and SAMtools. Bioinformatics. 2009;25: 2078–2079. doi:10.1093/bioinformatics/btp352

Broad Institute. Picard tools. https://broadinstitute.github.io/picard/. 2016; Available: http://broadinstitute.github.io/picard/

Schmidt S. (GATK) The Genome Analysis Toolkit: A MapReduce framework for analyzing next-generation DNA sequencing data. Proceedings of the International Conference on Intellectual Capital, Knowledge Management & Organizational Learning. 2009;20: 254–260. doi:10.1101/gr.107524.110.20

Purcell S , Neale B , Todd-Brown K , Thomas L , Ferreira MAR , Bender D , et al. PLINK: a tool set for whole-genome association and population-based linkage analyses. Am J Hum Genet. 2007;81: 559–575. doi:10.1086/519795

Chang CC , Chow CC , Tellier LC , Vattikuti S , Purcell SM , Lee JJ. Second-generation PLINK: rising to the challenge of larger and richer datasets. Gigascience. 2015;4: 7.

Patterson N , Price AL , Reich D. Population Structure and Eigenanalysis. PLoS Genetics. 2006;2: e190. doi:10.1371/journal.pgen.0020190

Skoglund P , Mallick S , Bortolini MC , Chennagiri N , Hünemeier T , Petzl-Erler ML , et al. Genetic evidence for two founding populations of the Americas. Nature. 2015;525: 104–108.

Alexander DH , Novembre J , Lange K. Fast model-based estimation of ancestry in unrelated individuals. Genome Res. 2009;19: 1655–1664. doi:10.1101/gr.094052.109

Jakobsson M , Rosenberg NA. CLUMPP: a cluster matching and permutation program for dealing with label switching and multimodality in analysis of population structure. Bioinformatics. 2007;23: 1801–1806. doi:10.1093/bioinformatics/btm233

Reich D , Patterson N , Campbell D , Tandon A , Mazieres S , Ray N , et al. Reconstructing Native American population history. Nature. 2012;488: 370–374. doi:10.1038/nature11258

Moorjani P , Patterson N , Hirschhorn JN , Keinan A , Hao L , Atzmon G , et al. The history of african gene flow into Southern Europeans, Levantines, and Jews. PLoS Genetics. 2011;7. doi:10.1371/journal.pgen.1001373

Loh PR , Lipson M , Patterson N , Moorjani P , Pickrell JK , Reich D , et al. Inferring admixture histories of human populations using linkage disequilibrium. Genetics. 2013;193: 1233–1254. doi:10.1534/genetics.112.147330

Prüfer K , Racimo F , Patterson N , Jay F , Sankararaman S , Sawyer S , et al. The complete genome sequence of a Neanderthal from the Altai Mountains. Nature. 2014;505: 43–9. doi:10.1038/nature12886

Yi X , Liang Y , Huerta-Sanchez E , Jin X , Cuo ZXP , Pool JE , et al. Sequencing of 50 Human Exomes Reveals Adaptation to High Altitude. Science. 2010;329: 75–78. doi:10.1126/science.1190371

